# Glycopeptide antibiotic teicoplanin inhibits cell entry of SARS-CoV-2 by suppressing the proteolytic activity of cathepsin L

**DOI:** 10.1101/2020.02.05.935387

**Authors:** Fei Yu, Ting Pan, Feng Huang, Ruosu Ying, Jun Liu, Huimin Fan, Junsong Zhang, Weiwei Liu, Yingtong Lin, Yaochang Yuan, Tao Yang, Rong Li, Xu Zhang, Xi Lv, Qianyu Chen, Anqi Liang, Fan Zou, Bingfeng Liu, Fengyu Hu, Xiaoping Tang, Linghua Li, Kai Deng, Xin He, Hui Zhang, Yiwen Zhang, Xiancai Ma

## Abstract

Since the outbreak of the coronavirus disease 2019 (COVID-19) caused by severe acute respiratory syndrome coronavirus 2 (SARS-CoV-2), the public health worldwide has been greatly threatened. The development of an effective treatment for this infection is crucial and urgent but is hampered by the incomplete understanding of the viral infection mechanism and the lack of specific antiviral agents. We previously reported that teicoplanin, a glycopeptide antibiotic that has been commonly used in the clinic to treat bacterial infection, significantly restrained the cell entry of Ebola virus, SARS-CoV and MERS-CoV by specifically inhibiting the activity of cathepsin L (CTSL). Here, we found that the cleavage sites of CTSL on the Spike of SARS-CoV-2 were highly conserved among all the variants. The treatment with teicoplanin suppressed the proteolytic activity of CTSL on Spike and prevented the cellular infection of different pseudotyped SARS-CoV-2 viruses. Teicoplanin potently prevented the entry of authentic SARS-CoV-2 into the cellular cytoplasm with an IC_50_ of 2.038 μM for the Wuhan-Hu-1 reference strain and an IC_50_ of 2.116 μM for the SARS-CoV-2 (D614G) variant. The pre-treatment of teicoplanin also prevented SARS-CoV-2 infection in hACE2 mice. In summary, our data reveal that CTSL is required for both SARS-CoV-2 and SARS-CoV infection and demonstrate the therapeutic potential of teicoplanin for universal anti-CoVs intervention.

**Importance:** Disease prevention and treatment are two important countermeasures to end the coronavirus disease 2019 (COVID-19). However, severe acute respiratory syndrome coronavirus 2 (SARS-CoV-2), the causative agent of COVID-19, evolves all the time, resulting in the emerging of many epidemic SARS-CoV-2 mutants, which significantly impairs the effectiveness of early strain-based vaccines and antibodies. Developing universal vaccines and broad-spectrum antiviral drugs are essential to confront SARS-CoV-2 mutants including those may emerge in the future. Our study reported here showed that the cleavage sites of cellular cathepsin L (CTSL) are highly conserved among all the SARS-CoV-2 mutants and SARS-CoV. The CTSL inhibitor teicoplanin not only inhibited the cell entry of two live SARS-CoV-2 strains and various pseudotyped viruses but also prevented live virus infection in animal models. Based on our previous finding that teicoplanin also inhibited SARS-CoV and MERS-CoV infection, we believe that teicoplanin possesses the potential to become a universal anti-CoVs drug.

## Introduction

Coronaviruses (CoVs) are enveloped, positive sense single-stranded RNA viruses (1, 2). Many members of the coronavirus family are life-threatening human pathogens and can cause severe respiratory diseases, such as severe acute respiratory syndrome-associated coronavirus (SARS-CoV) emerged in 2003 and middle east respiratory syndrome coronavirus (MERS-CoV) emerged in 2012 (3-9). Since December 2019, a novel coronavirus has emerged and spread globally, resulting in millions of pneumonia cases around the world (10-14). This novel coronavirus, named SARS-CoV-2, belongs to the beta-coronavirus according to the sequence released (13, 15). Evolutionary analyses have shown that SARS-CoV-2 shares 79% homology with SARS-CoV and 50% homology with MERS-CoV (15-17). During the last two years, many independent dominant SARS-CoV-2 variants have emerged locally and circulated globally, which included B.1.1.7 (Alpha), B.1.351 (Beta), P.1 (Gamma), B.1.429 (Epsilon), B.1.525 (Eta), B.1.526 (Iota), B.1.617.1 (Kappa), B.1.617.2 (Delta), B.1.621 (Mu) and C.37 (Lambda) (18-26). Given the high frequency of mutation, the high infectious rate and the lack of effective treatment for SARS-CoV-2, it is urgent to develop an efficient antiviral drug for SARS-CoV-2 and its mutants.

The Spike (S) glycoproteins, which cover on the surface of virions, mediate the viral entry into host cells and determine the host range of coronaviruses (27-29). The infection of both SARS-CoV and SARS-CoV-2 is initiated by the attachment of the S protein to the host receptor angiotensin-converting enzyme 2 (ACE2) (29-31), followed by the S protein priming by cellular proteases such as TMPRSS2 (31-34). The viruses are then transported into host cells through the early endosomes and late endosomes, and subsequently endo / lysosomes. For SARS-CoV, the primed S proteins are further cleaved by other proteases such as cysteine proteinase cathepsin L (CTSL) within endocytic vesicles to complete the activation (35-38). The activated S proteins then mediate the fusion of viral and cellular membranes, resulting in the release of SARS-CoV genome into the cytoplasm.

We previously found that teicoplanin, a commonly used clinical glycopeptide antibiotic, potently suppressed the cellular entry of Ebola virus, SARS-CoV, and MERS-CoV (39). Further mechanism investigation revealed that teicoplanin blocked the virus entry by specifically inhibiting the proteolytic activity of CTSL, indicating the potential of teicoplanin as an effective drug for CTSL-dependent viral infection. In this study, we investigated the role of CTSL in SARS-CoV-2 entry and tested the inhibitory effect of teicoplanin and homologs on the viral entry process. We found that the cleavage sites of CTSL were highly conserved among the S sequences of various epidemic SARS-CoV-2 mutants and SARS-CoV. The loss of CTSL significantly crippled SARS-CoV-2 infection, while the overexpression of CTSL significantly increased the infectivity of SARS-CoV-2. Meanwhile, teicoplanin and dalbavancin, but not vancomycin, exhibited remarkable inhibitory activity toward the entry of SARS-CoV-2. Teicoplanin was able to inhibit the entry of all the major epidemic SARS-CoV-2 mutants. Further mechanism study indicated that teicoplanin inhibited SARS-CoV-2 entry by inhibiting the proteolytic activity of CTSL on S proteins. More importantly, teicoplanin inhibited the entry of authentic SARS-CoV-2 viruses with an IC_50_ lower than 5 μM (2.038 μM for the original strain, 2.116 μM for the D614G variant). The pre-treatment of teicoplanin also prevented the infection of authentic SARS-CoV-2 in mice models. Combined with our previous finding that teicoplanin inhibited the entry of SARS-CoV and MERS-CoV, our study reported here indicated that the CTSL inhibitor teicoplanin could be a universal anti-CoVs drug.

## Results

### SARS-CoV-2 infection depended on the activity of CTSL

The proteolytic processing of the S protein is essential for SARS-CoV entry and fusion. Many host proteases, including TMPRSS2 and CTSL, are involved in the priming and activation of the SARS-CoV S protein, and some of which also have been identified and experimentally validated in SARS-CoV-2 infection (31, 32, 40-45). To systematically identify cellular proteases and receptors which mediated the entry and fusion of SARS-CoV-2 to target cells, we knocked down ten major proteases and receptors in HEK293T cells, which included CTSL, CTSB, CTSK, TMPRSS2, TMPRSS11A, TMPRSS11D, Furin, PLG, DPP4 and ACE2. Subsequently, these cells were infected by pseudotyped SARS-CoV-2 S / HIV-1 viruses which harbored an integrated *luciferase* gene. The expression of luciferase indicated the entry and expression of pseudotyped virus. We found that the absence of CTSL, TMPRSS2, Furin or ACE2 significantly decreased the pseudotyped SARS-CoV-2 virus infection (**Fig. 1A**). The ACE2 protein has been identified as the major receptor of SARS-CoV-2 (17). Both TMPRSS2 and Furin also have been found to be essential for efficient infection of SARS-CoV-2 (31, 42). To determine whether CTSL is also involved in SARS-CoV-2 S protein activation, we compared the cleavage sites of CTSL in the gene sequences encoding the SARS-CoV and SARS-CoV-2 S proteins. After alignment, we found that the cleavage sites of CTSL were well-conserved between SARS-CoV and SARS-CoV-2 S proteins (**Fig. 1B**). Moreover, the cleavage sites of CTSL on S proteins were also highly conserved among all the major epidemic SARS-CoV-2 variants including D614 (Wuhan-Hu-1), G614 (SYSU-IHV), B.1.1.7 (Alpha), B.1.351 (Beta), P.1 (Gamma), B.1.429 (Epsilon), B.1.525 (Eta), B.1.526 (Iota), B.1.617.1 (Kappa), B.1.617.2 (Delta), B.1.621 (Mu) and C.37 (Lambda) (18-26) (**Fig. 1C**). Previously, CTSL has been found to play pivotal roles in SARS-CoV infection by cleaving and activating S proteins (38, 41, 46). We speculated that CTSL might also participate in SARS-CoV-2 entry and fusion. Thus, we overexpressed CTSL proteins in HEK293T cells which were subsequently infected by pseudotyped SARS-CoV-2 S / HIV-1 viruses. We found that the infectivity of pseudotyped virus to HEK293T cells was linearly and positively correlated with the expression level of CTSL (**Fig. 1D**). We also co-overexpressed ACE2 with CTSL in HEK293T cells. We found that the co-overexpression of CTSL significantly increased the infectivity of pseudotyped virus compared with ACE2-overexpression only (**Fig. 1E**). These results indicated that the effective infection of SARS-CoV-2 to host cells depended on the proteolytic activity of CTSL.

**Figure 1.**
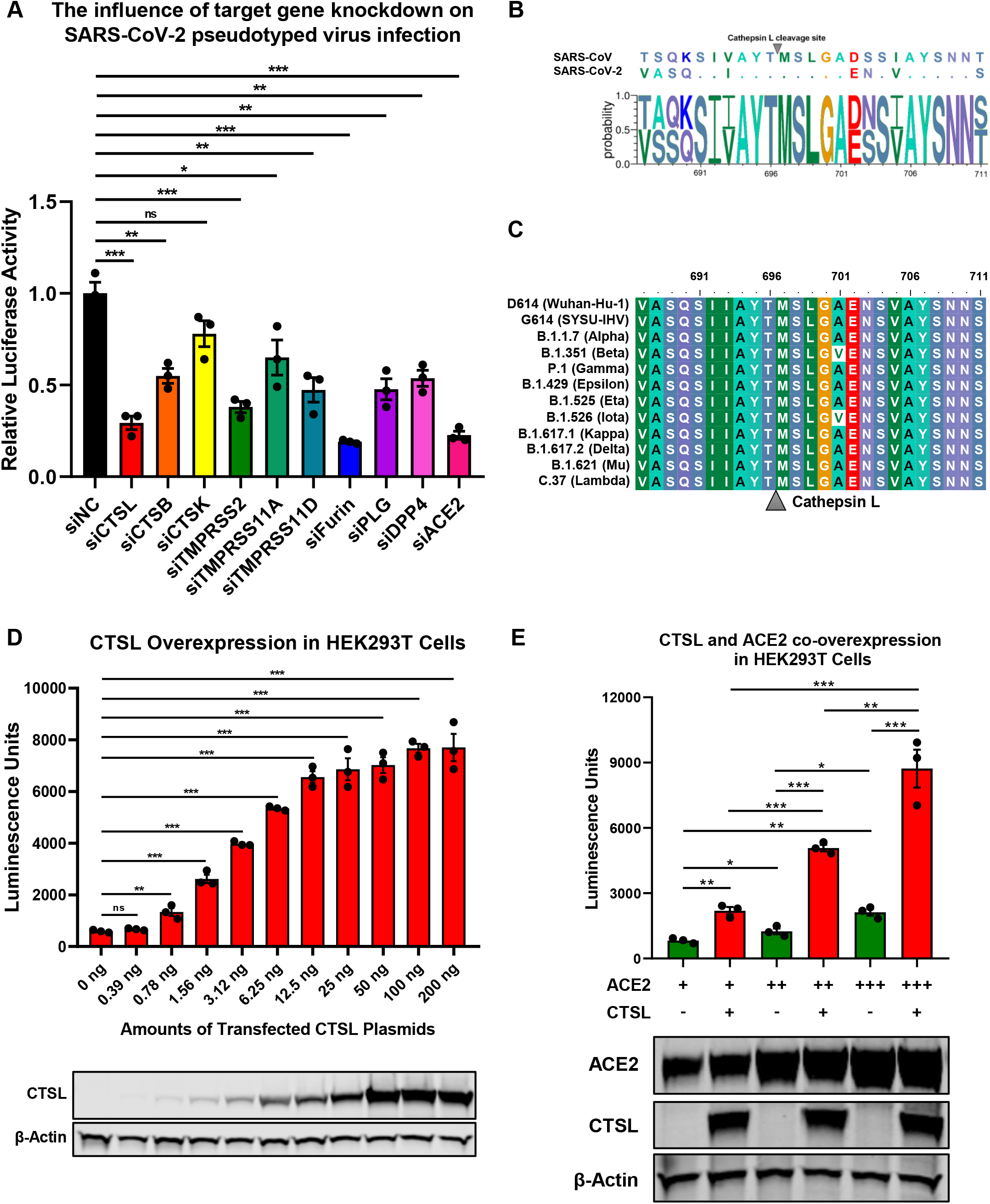
SARS-CoV-2 infection depended on the activity of CTSL. (**A**) CTSL, CTSB, CTSK, TMPRSS2, TMPRSS11A, TMPRSS11D, Furin, PLG, DPP4 and ACE2 in HEK293T cells were knocked down by siRNAs. These cells were infected with pseudotyped SARS-CoV-2 viruses 24 hours post transfection. The intracellular luciferase activity was measured after another 24 hours. The fold change of luciferase expression in each group was normalized to si-negative control (siNC) group (n=3). (**B**) Sequence alignment based on the consensus S protein sequences of SARS-CoV and SARS-CoV-2. The overall height of the stack indicated the sequence conservation at that position, while the height of symbols within the stack indicated the relative frequency (Y-axis) of each amino acid at that position (X-axis). (**C**) The multiple alignments were created based on the region containing the cleavage site of cathepsin L (CTSL) (SIIAYTMSLGA) in the S protein of SARS-CoV-2. The S proteins of 12 different SARS-CoV-2 variants including D614 (Wuhan-Hu-1), G614 (SYSU-IHV), B.1.1.7 (Alpha), B.1.351 (Beta), P.1 (Gamma), B.1.429 (Epsilon), B.1.525 (Eta), B.1.526 (Iota), B.1.617.1 (Kappa), B.1.617.2 (Delta), B.1.621 (Mu) and C.37 (Lambda) were involved. The identity/similarity shading with the color refers to the chemistry of each amino acid at that position. (**D**) HEK293T cells in 96-well plate were transfected with two-fold serially diluted CTSL-expressing plasmids, ranging from 0.39 ng to 200 ng. Cells were infected with pseudotyped SARS-CoV-2 S /HIV-1 viruses 24 hours post transfection. Another 48 hours post infection, cells were lysed and measured for the amounts of luciferase which were represented by luminescence units (n=3). Western blot with antibodies against CTSL was conducted to confirm the expression of CTSL plasmids. β-Actin was immunoblotted as internal control. (**E**) HEK293T cells were transfected with different amounts of ACE2-expressing plasmids. Another groups of cells were co-transfected with CTSL-expressing plasmids. These cells were infected with pseudotyped SARS-CoV-2 S / HIV-1 viruses 24 hours post transfection. The amounts of luciferase within each group were measured 48 hours post infection and represented as luminescence units (n=3). The expression of ACE2 and CTSL was confirmed by western blot. Data in (**A**) and (**D-E**) represented as mean ± SEM in triplicate. P-values in (**A**) and (**D**) were calculated by one-way ANOVA with Dunnett’s multiple comparison test which compared the mean of each group with the mean of the control group. P-values in (**E**) were calculated by one-way ANOVA with Tukey’s multiple comparison test which compared the mean of each group with the mean of every other group. ns = p ≥ 0.05, *p < 0.05, **p < 0.01, ***p < 0.001.

### Teicoplanin specifically inhibited the entry of SARS-CoV-2

Previously, we have found that teicoplanin, a glycopeptide antibiotic which inhibited CTSL activity, suppressed the entry of SARS-CoV, MERS-CoV and Ebola viruses (39). We speculated that teicoplanin might also be able to block the entry of SARS-CoV-2. Thus, we conducted the pseudotyped virus entry upon drug treatment assay. We generated a highly sensitive HEK293T-hACE2 cell line which constitutively expressed high level of hACE2 receptors. HEK293T-hACE2 cells were co-incubated with teicoplanin and pseudotyped SARS-CoV-2 S / HIV-1 virus. The infectivity of pseudotyped virus, which was represented by the amounts of luciferase within HEK293T-hACE2 cells, was measured 48 hours post infection (**Fig. 2A**). To exclude the possibility that teicoplanin inhibited the early events of the pseudotyped HIV-1 life cycle, pseudotyped VSV-G / HIV-1 viruses bearing vesicular stomatitis virus (VSV) glycoproteins were also packaged and treated as the negative control. Pseudotyped SARS-CoV S / HIV-1 viruses bearing SARS-CoV S were packaged and treated as the positive control. The results showed that teicoplanin effectively inhibited the entry of both SARS-CoV-2 and SARS-CoV pseudotyped viruses in a dose-dependent manner, whereas teicoplanin treatment did not affect the infection of pseudotyped VSV-G / HIV-1 viruses (**Fig. 2B and 2C**).

**Figure 2.**
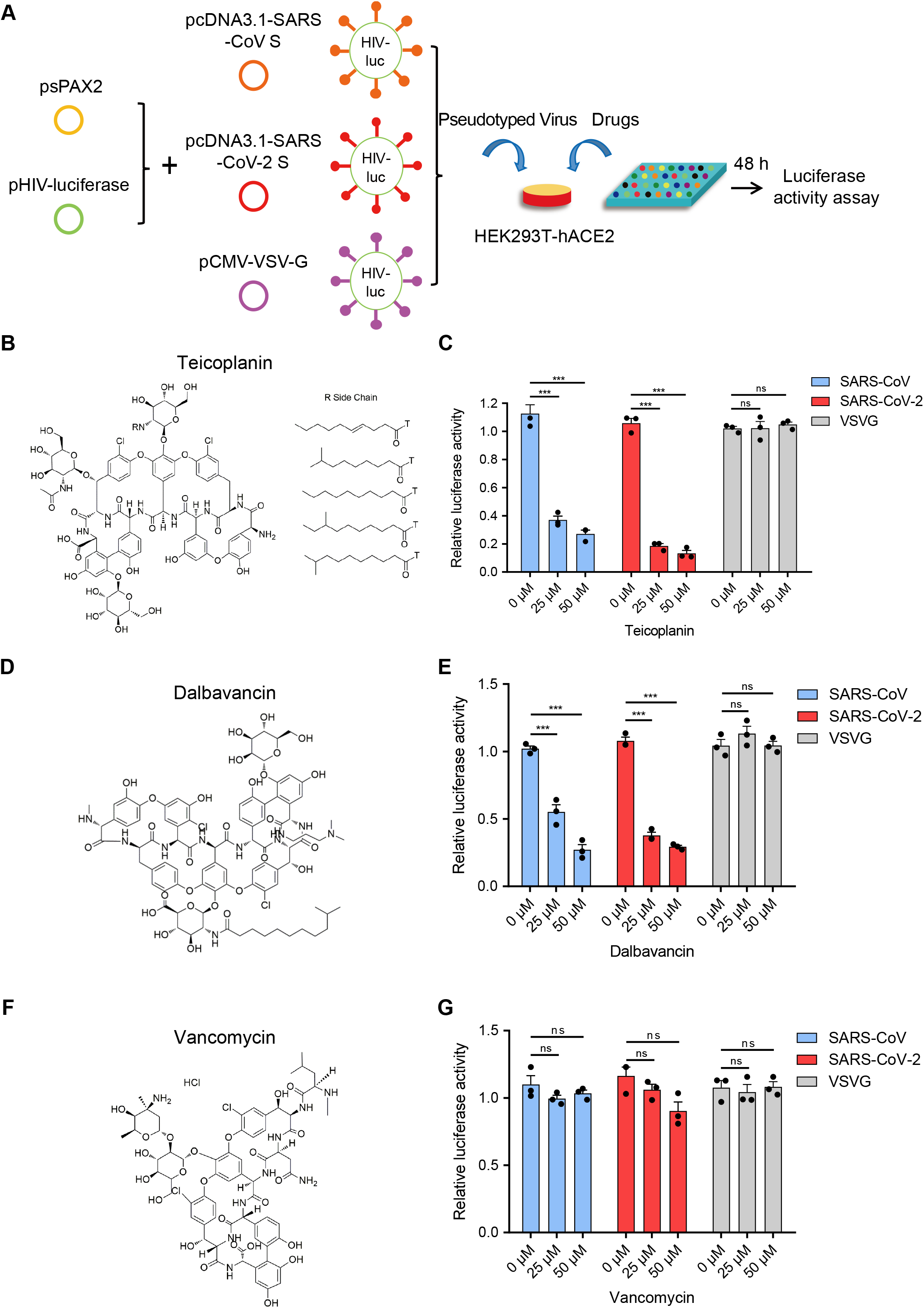
Teicoplanin specifically inhibited the entry of SARS-CoV-2. (**A**) Schematic of the pseudotyped virus entry upon drug treatment assay. To package different pseudotyped viruses, psPAX2 plasmids and pHIV-Luciferase plasmids were co-transfected into HEK293T cells with pcDNA3.1-SARS-CoV S, pcDNA3.1-SARS-CoV-2 S and pCMV-VSV-G plasmids respectively. HEK293T-hACE2 cells were incubated with drugs and different pseudotyped viruses. The amounts of luciferase within cells were measured 48 hours post infection. (**B**) Chemical structure of teicoplanin. (**C**) HEK293T-hACE2 cells were treated with 0 μM, 25 μM and 50 μM teicoplanin respectively, followed by infecting with different pseudotyped viruses including SARS-CoV S / HIV-1, SARS-CoV-2 S / HIV-1 and VSV-G / HIV-1. The intracellular luciferase activity was measured after 48 hours post infection (n=3). (**D**) Chemical structure of dalbavancin. (**E**) HEK293T-hACE2 cells were treated as in (**C**), except that the drug was replaced with dalbavancin (n=3). (**F**) Chemical structure of vancomycin. (**G**) HEK293T-hACE2 cells were treated as in (**C**), except that the drug was replaced with vancomycin (n=3). Data in (**C**), (**E**) and (**G**) represented as mean ± SEM in triplicate. P-values were calculated by two-way ANOVA with Dunnett’s multiple comparisons test which compared the mean of each group with the mean of the control group. ns = p ≥ 0.05, ***p < 0.001.

Teicoplanin homologs including dalbavancin also have specific inhibitory effects on CTSL based on our previous study (39). While vancomycin, another glycopeptide antibiotic which was clinically used for Gram-positive bacterial infections, did not show inhibitory activity on CTSL. Therefore, we further tested whether dalbavancin and vancomycin could inhibit the entry of SARS-CoV-2. Similar to teicoplanin, dalbavancin effectively inhibited both SARS-CoV-2 and SARS-CoV pseudotyped viruses entering into HEK293T-hACE2 cells in a dose-dependent manner, but it did not affect pseudotyped VSV-G / HIV-1 viruses infection (**Fig. 2D and 2E**). In contrast, vancomycin did not show any inhibitory activity on the infection of pseudotyped SARS-CoV-2, SARS-CoV or VSV-G viruses (**Fig. 2F and 2G**). Taken together, these results indicated that CTSL inhibitors teicoplanin and its homolog dalbavancin could suppress the entry of SARS-CoV-2.

### Teicoplanin inhibited the entry of SARS-CoV-2 by inhibiting the activity of CTSL

To further confirm that teicoplanin inhibited the entry of SARS-CoV-2, we investigated the antiviral activity of teicoplanin on authentic (live) SARS-CoV-2. We obtained two authentic SARS-CoV-2 strains. One was SARS-CoV-2 D614 virus (Wuhan-Hu-1) which was provided by Guangdong Provincial Center for Disease Control and Prevention (GDCDC). The other was SARS-CoV-2 G614 virus (SYSU-IHV) which was isolated by us from the sputum sample of an infected patient (47). We found that teicoplanin effectively inhibited the entry of both authentic strains with a half maximal inhibitory concentration (IC_50_) of 2.038 μM for the Wuhan-Hu-1 reference strain and an IC_50_ of 2.116 μM for the SARS-CoV-2 (D614G) variant (**Fig. 3A and 3B**). Given that the serum concentrations of teicoplanin in patients are at least 15 mg / L (8.78 µM) after the loading dose treatment for most Gram-positive bacterial infections, our data indicated that teicoplanin was able to potently suppress the entry of SARS-CoV-2 of both the original Wuhan-Hu-1 strain and the D614G mutation strain at a relatively low and safe dose (39).

**Figure 3.**
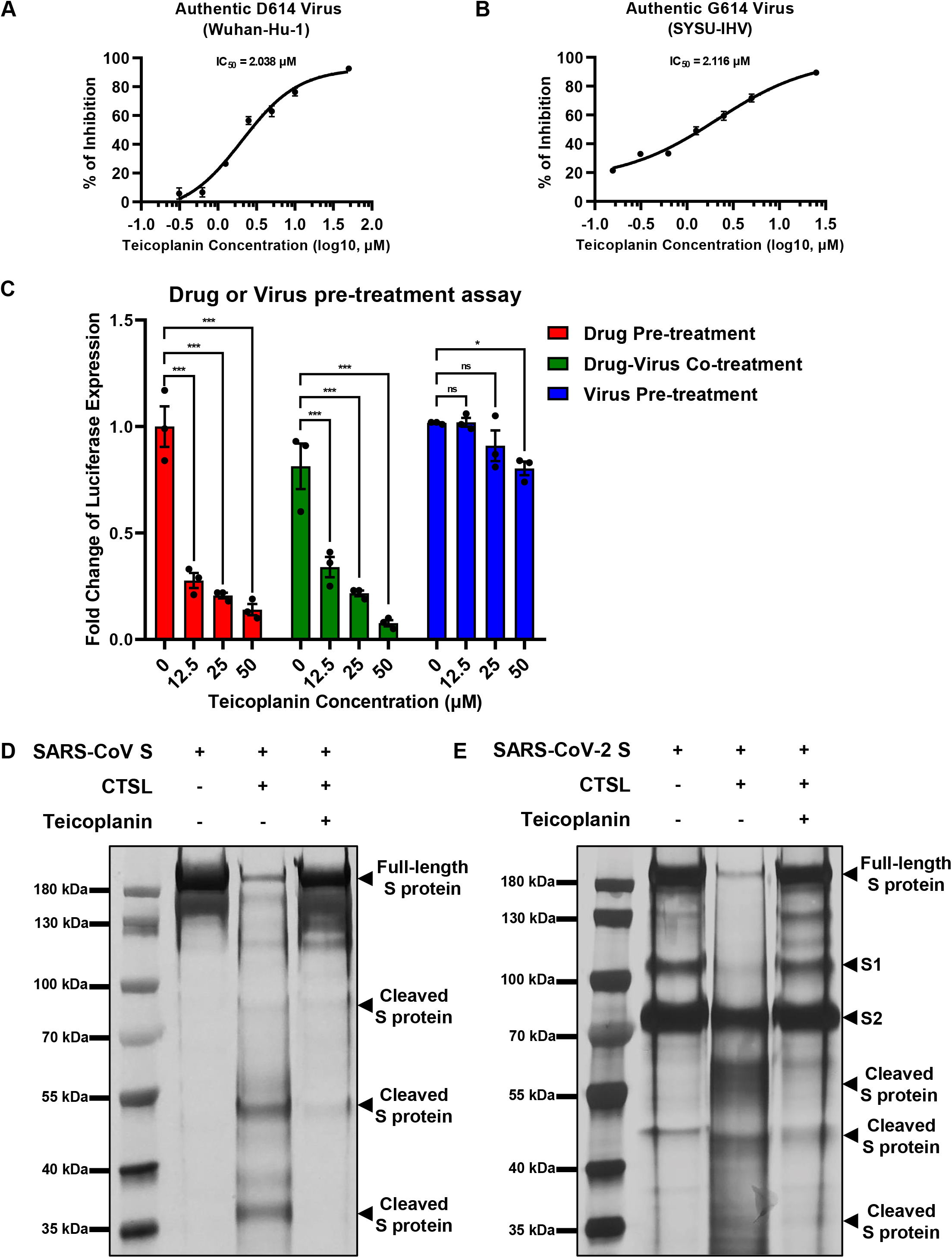
Teicoplanin inhibited the entry of SARS-CoV-2 by inhibiting the activity of CTSL. (**A**) HEK293T-hACE2 cells were co-incubated with authentic SARS-CoV-2 D614 (Wuhan-Hu-1) virus and two-fold serially diluted teicoplanin. At 48 hours post incubation, the supernatant in each group was collected and proceeded to RNA extraction. Viral RNA copies in supernatant were quantified by one-step SARS-CoV-2 RNA detection kit. The IC_50_ of teicoplanin against SARS-CoV-2 D614 virus was calculated according to these viral RNA copies within each group (n=3). (**B**) The IC_50_ of teicoplanin against authentic SARS-CoV-2 G614 (SYSU-IHV) virus was determined as in (**A**) (n=3). (**C**) In the first group, HEK293T-hACE2 cells were pre-treated with 0 μM, 12.5 μM, 25 μM and 50 μM teicoplanin respectively. Four hours later, cells were infected with pseudotyped SARS-CoV-2 S / HIV-1 virus. In the second group, cells were co-treated with different concentrations of teicoplanin and pseudotyped SARS-CoV-2 S / HIV-1 virus simultaneously. In the third group, cells were pre-infected with pseudotyped SARS-CoV-2 S / HIV-1 virus. Four hours post infection, cells were treated with different concentrations of teicoplanin. The amounts of luciferase within cells were quantified 48 hours post infection. The fold changes of luciferase expression within each sample were calculated by normalizing to those in cells treated with 0 μM teicoplanin (n=3). (**D-E**) The *in vitro* purified 250 ng CTSL proteins were added into CTSL assay buffer for activation. In another group, CTSL were also co-incubated with 50 μM teicoplanin. After activation, the *in vitro* purified SARS-CoV S or SARS-CoV-2 S proteins were added into each group. S protein only group was set as control group. The digested proteins were proceeded to SDS-PAGE and analyzed by silver staining. Data in (**A-C**) represented as mean ± SEM in triplicate. Inhibition curves in (**A-B**) were generated by log (inhibitor) vs. response nonlinear fit. P-values in (**C**) were calculated by two-way ANOVA with Dunnett’s multiple comparisons test which compared the mean of each group with the mean of the control group. ns = p ≥ 0.05, *p < 0.05, ***p < 0.001.

To elucidate whether the target of teicoplanin was the virus itself, or the host cell, or both, we conducted drug / virus pre-treatment assay. In the first group, HEK293T-hACE2 cells were pre-treated with different concentrations of teicoplanin followed by infecting with pseudotyped SARS-CoV-2 (drug pre-treatment group). In the second group, cells were co-incubated with both teicoplanin and pseudotyped virus (drug-virus co-treatment group). In the third group, cells were pre-infected with pseudotyped virus followed by treating with teicoplanin (virus pre-treatment group). We found that the infectivity of pseudotyped SARS-CoV-2 in both drug pre-treatment group and drug-virus co-treatment group was negatively correlated with the concentration of teicoplanin, whereas the infectivity of pseudotyped virus in virus pre-treatment group was almost unchanged upon the treatment of different concentrations of teicoplanin (**Fig. 3C**). These results indicated that teicoplanin targeted host cells rather than viral particles.

Our previous report has revealed that teicoplanin targets on CTSL directly within host cells (39). To provide direct evidence that teicoplanin inhibiting SARS-CoV-2 entry via inhibiting the activity of CTSL as well, we conducted *in vitro* CTSL enzymatic inhibition assay. The *in vitro* purified CTSL proteins were firstly activated in CTSL assay buffer, followed by incubating with SARS-CoV S or SARS-CoV-2 S. In another group, pre-activated CTSL proteins were co-incubated with different S proteins and teicoplanin. Then the CTSL- and teicoplanin-treated S proteins were proceeded to SDS-PAGE and silver staining. We found that CTSL proteins were able to effectively cleave both SARS-CoV S and SARS-CoV-2 S (**Fig. 3D and 3E**). However, the co-treatment of teicoplanin with CTSL proteins inhibited the enzymatic activity of CTSL on both S proteins, resulting in the presence of more full-length S proteins (**Fig. 3D and 3E**). Our above results further confirmed that teicoplanin inhibited the entry of SARS-CoV-2 by directly inhibiting the proteolytic activity of CTSL within host cells.

### Teicoplanin inhibited the entry of various SARS-CoV-2 mutants

Since December 2019, many SARS-CoV-2 mutants have emerged locally and spread worldwide, such as B.1.1.7 (Alpha), B.1.351 (Beta), P.1 (Gamma), B.1.429 (Epsilon), B.1.525 (Eta), B.1.526 (Iota), B.1.617.1 (Kappa), B.1.617.2 (Delta), B.1.621 (Mu) and C.37 (Lambda) (18-26). We have found that the cleavage sites of CTSL on S proteins of these mutants were highly conserved (**Fig. 1C**). Thus, we speculated that teicoplanin-mediated inhibition of CTSL activity might also cripple the cell entry of SARS-CoV-2 mutants. To evaluate whether teicoplanin still was able to inhibit the infection of these variants, we constructed different S-expressing plasmids which were derived from various SARS-CoV-2 mutants (**Fig. 4A**). Similar to the package of pseudotyped SARS-CoV-2 (D614) S / HIV-1 viruses, we packaged ten different pseudotyped SARS-CoV-2 viruses based on the above S mutants. HEK293T-hACE2 cells were co-incubated with different pseudotyped viruses and two-fold serially diluted teicoplanin, followed by the measurement of the luciferase activity which could represent viral infectivity. The IC_50_ of teicoplanin against different pseudotyped viruses was calculated based on the percentages of viral inhibition. We found that all the IC_50_ of teicoplanin against the entry of these viruses were below 5 μM (3.002 μM for B.1.351 / Beta, 3.117 μM for P.1 / Gamma, 3.056 μM for B.1.429 / Epsilon, 2.041 μM for B.1.525 / Eta, 1.963 μM for B.1.526 / Iota, 2.188 μM for B.1.617.1 / Kappa, 2.300 μM for B.1.617.2 / Delta, 1.998 μM for B.1.621 / Mu, 2.306 μM for C.37 / Lambda), except that the IC_50_ of teicoplanin against pseudotyped SARS-CoV-2 (B.1.1.7 / Alpha) S / HIV-1 viruses was 5.423 μM (**Fig. 4B-4K**). Taken together, our above results indicated that the CTSL inhibitor teicoplanin was able to inhibit the entry of different SARS-CoV-2 variants.

**Figure 4.**
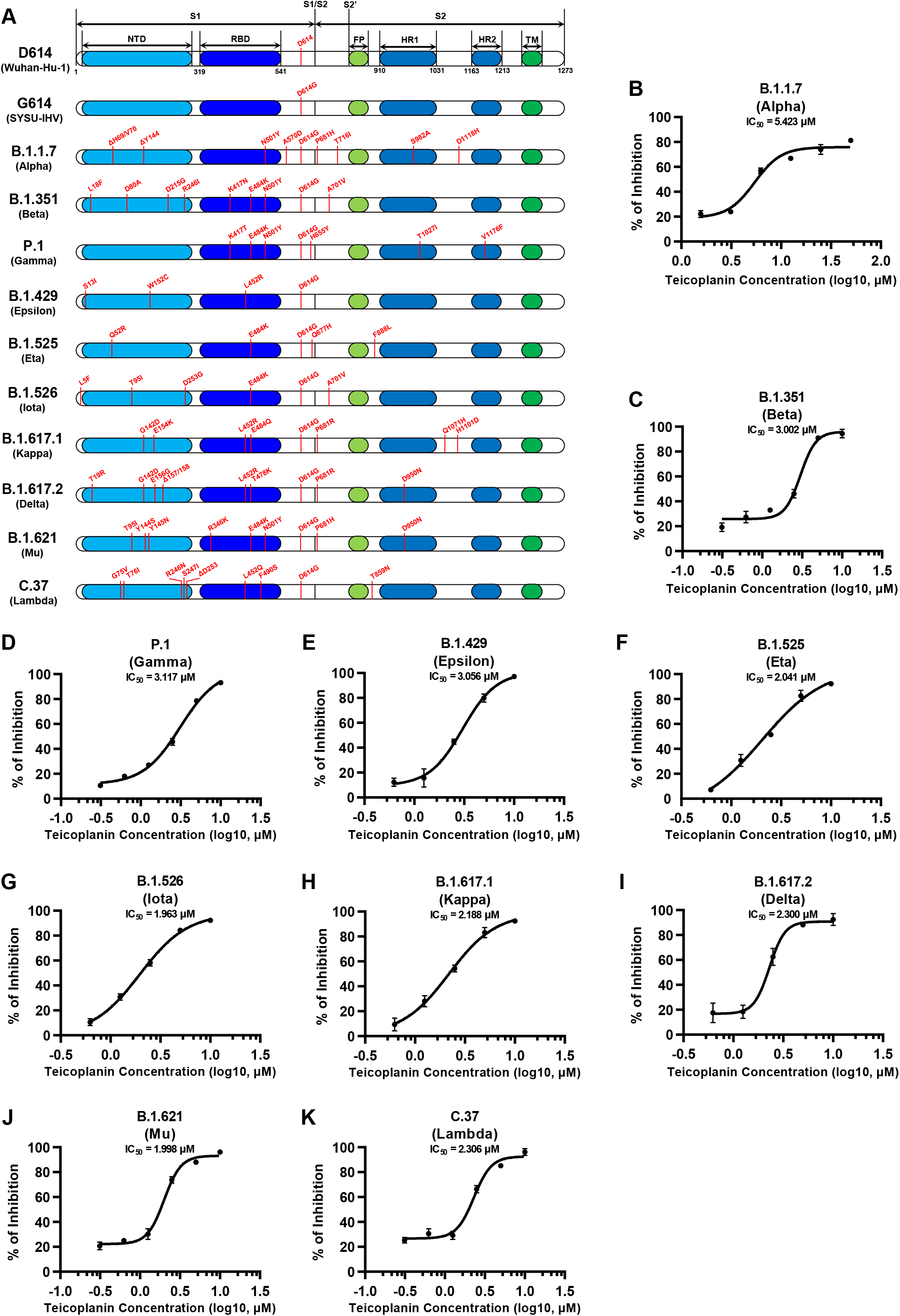
Teicoplanin inhibited the entry of various SARS-CoV-2 mutants. (**A**) Schematics of the Spike proteins of 12 different SARS-CoV-2 mutants which included D614 (Wuhan-Hu-1), G614 (SYSU-IHV), B.1.1.7 (Alpha), B.1.351 (Beta), P.1 (Gamma), B.1.429 (Epsilon), B.1.525 (Eta), B.1.526 (Iota), B.1.617.1 (Kappa), B.1.617.2 (Delta), B.1.621 (Mu) and C.37 (Lambda). The mutation sites were shown alongside each backbone and indicated in red. (**B-K**) HEK293T-hACE2 cells were co-incubated with different pseudotyped SARS-CoV-2 S / HIV-1 viruses and serially diluted teicoplanin. The amounts of luciferase within each group were measured 48 hours post infection and represented as luminescence units. The IC_50_ of teicoplanin against these pseudotyped SARS-CoV-2 mutants was calculated based on the amounts of luciferase within each group (n=3). Data in (**B-K**) represented as mean ± SEM in triplicate. Inhibition curves were generated by log (inhibitor) vs. response nonlinear fit.

### Teicoplanin prevented SARS-CoV-2 infection in hACE2 mice

To evaluate whether the treatment of teicoplanin could protect individuals from SARS-CoV-2 infection, we conducted mice infection experiments upon teicoplanin treatment. We utilized K18-hACE2 mice which were generated by knocking in the human K18 promoter-driven human ACE2 within the mouse Hipp11 (H11) “safe-harbor” locus. hACE2 mice were intraperitoneally administrated with 100 mg / kg body weight teicoplanin or equal volume of saline, followed by intranasally challenging with 1×10^5^ focus-forming units (FFU) of authentic SARS-CoV-2 D614 virus (n=4 in each group). These mice were euthanized 5 days post infection (**Fig. 5A**). Lung tissues of each mice were proceeded to SARS-CoV-2 viral RNA quantification, hematoxylin & eosin (HE) and immunohistochemistry (IHC) analysis. We found that lung tissues of hACE2 mice in saline group harbored large amounts of viral RNA copies (1.9×10^4^, 2.3×10^5^, 1.2×10^5^, and 8.8×10^4^ copies per ml) (**Fig. 5B**). While lung tissues of hACE2 mice in teicoplanin group harbored only few numbers of viral RNA copies (less than 10 copies per ml in average) (**Fig. 5B**). HE and IHC assays also revealed that lung tissues in mice from saline group were severely damaged upon SARS-CoV-2 challenge, which were interspersed with thickened alveolar septa, collapsed alveoli and Nucleoprotein (N) protein-expressing cells (**Fig. 5C**). Whereas no pathological changes and N-expressing cells were observed in teicoplanin treatment group (**Fig. 5C**). These results indicated that teicoplanin treatment was able to prevent the infection of authentic SARS-CoV-2 virus in hACE2 mice.

**Figure 5.**
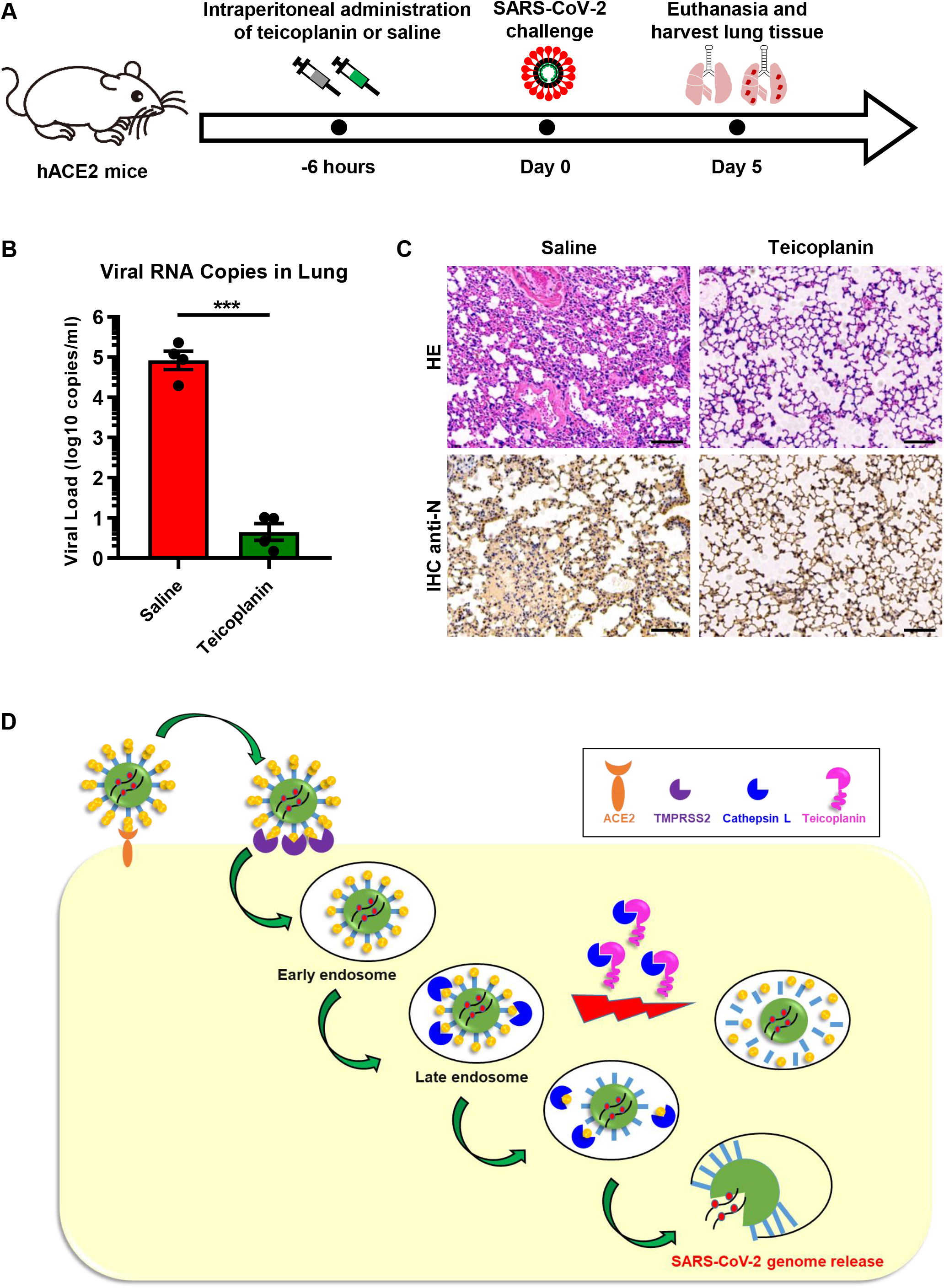
Teicoplanin prevented SARS-CoV-2 infection in hACE2 mice. (**A**) Schematic of mice experiment procedure. hACE2 mice were intraperitoneally administrated with 100 mg/kg body weight teicoplanin or saline (n=4 in each group). Six hours later, each mice was challenged with 1×10^5^ FFU of authentic SARS-CoV-2 D614 viruses. Another 5 days later, mice were euthanized to harvest lung tissues which were proceeded to viral RNA detection, HE and IHC. (**B**) Viral RNA copies in lung tissues of virus-challenged mice were quantified by one-step SARS-CoV-2 RNA detection kit and plotted as log10 copies per ml (n=4). (**C**) Lung tissues of mice from saline group and teicoplanin group were proceeded to HE staining and IHC with antibodies against N proteins. (**D**) Schematic of teicoplanin inhibiting the entry of SARS-CoV-2. The SARS-CoV-2 virus binds to the cellular receptor ACE2 by its Spike (S) proteins which cover the surface of the virus. The S-ACE2 binding event initiates the proteolytic process of TMPRSS2 to S protein on the cellular membrane, followed by the entry of virus to the early endosome. Then, the virion is transported into the late endosome where the S protein is further activated by cleavage with cysteine proteinase CTSL. After S activation by CTSL, the viral genome is released to the cytoplasm where viruses replicate and assemble. Teicoplanin can effectively block the proteolytic activity of CTSL, rendering the S protein unable to be activated. Without S activation, The SARS-CoV-2 virus is dissolved in the endosome gradually. Scale bars in (**C**) represented 100 μm. Data in (**B**) represented as mean ± SEM in quadruplicate. P-value was calculated by Student’s *t* test. ***p < 0.001.

## Discussion

To date, many drugs have been tested for treatment of COVID-19. Remdesivir showed some efficacy in COVID-19 patients, but many severe side effects were observed (48). Among patients hospitalized in metropolitan New York with COVID-19, treatment with hydroxychloroquine and / or azithromycin failed to significantly improve in-hospital mortality (49). Recently, Merck Sharp and Dohme (MSD) and its partner Ridgeback Biotherapeutics reported that the antiviral drug molnupiravir (MK-4482, EIDD-2801) reduced the risk of hospitalization or death by 50% compared to placebo for patients with mild or moderate COVID-19 based on their Phase III study (NCT04575597). Although previous reports also showed that molnupiravir could prevent SARS-CoV-2 infection and transmission in animal models, more clinical data of long-term monitoring need to be collected to investigate its potential side effects in the future clinical trials (50, 51). Specific treatment for COVID-19 is still lacking and urgently needed. Host cell entry is the first step of the viral life cycle and is an ideal process to develop potential drugs. In this study, we identified that teicoplanin could inhibit the entry of SARS-CoV-2 with an IC_50_ of lower than 2.5 µM. Teicoplanin not only exhibited remarkable inhibitory activity on various pseudotyped SARS-CoV-2 S / HIV-1 mutants entry, but also potently restrained the infection of authentic SARS-CoV-2 viruses including the original D614 reference strain and the later G614 variant. This inhibitory effect was also confirmed by animal experiment. Therefore, these findings may provide a novel therapeutic treatment to improve current antiviral therapy.

During the invasion phase, SARS-CoV-2 firstly binds to its receptor hACE2 on the surface of host cells. The interaction between the receptor-binding domain (RBD) of the S protein and hACE2 triggers conformational changes within the S protein, which renders the S protein susceptible to be activated by the host cell protease TMPRSS2 (31, 32, 40). Subsequently, the SARS-CoV-2 virus enters to the early endosome of the host cell through endocytosis or macropinocytosis. During the maturation process of the early endosome, the endosome gradually acidifies, which facilitates the entry of viruses into cells. The antiviral drug chloroquine, which can increase the endosomal pH to block virus infection, has been found to inhibit the entry of both SARS-CoV and SARS-CoV-2 (17, 52, 53). During the entry and fusion of SARS-CoV, the cysteine proteinase CTSL within the late endosome can further cleave the S protein and activate the membrane fusion, resulting in the release of viral genome (37, 38).

Several CRIPSR-mediated knock-out and animal infection experiments have highlighted the importance of CTSL in SARS-CoV-2 infection (43-45). Here we showed that knocking down CTSL potently inhibited SARS-CoV-2 entry. The overexpression of CTSL significantly increased the infectivity of pseudotyped SARS-CoV-2 viruses. Moreover, our data also demonstrated that teicoplanin was able to potently inhibit the infection of both authentic SARS-CoV-2 viruses and different pseudotyped SARS-CoV-2 mutants by inhibiting the enzymatic activity of CTSL and preventing S proteins further activation. Without complete activation of S proteins, the SARS-CoV-2 virus was gradually degraded within the endosome (**Fig. 5D**). Based on all these findings, we believe that the endosomal proteinase CTSL plays vital roles in the infection of SARS-CoV, MERS-CoV, SARS-CoV-2, and possibly other coronaviruses. Therefore, CTSL and its inhibitor teicoplanin provide important therapeutic potential for developing universal anti-CoVs intervention.

Teicoplanin is a glycopeptide antibiotic which is mainly used for serious infection caused by Gram-positive bacteria such as *Staphylococcus aureus* and *Streptococcus* (54-56). As a commonly used clinical antibiotic, teicoplanin is well known for its low toxicity, mild side effects, long half-life in blood plasma, convenient administration, and high safety. Clinically, the serum concentration of teicoplanin is at least 15 mg / L (8.78 µM) after the completion of the loading dose treatment for most Gram-positive bacterial infections (39). In this study, we found that teicoplanin, and its homolog dalbavancin, could inhibit the entry of SARS-CoV-2 in HEK293T-hACE2 cells expressing the key receptor ACE2. Importantly, teicoplanin was able to inhibit authentic SARS-CoV-2 viruses entry with an IC_50_ of 2.038 μM for the Wuhan-Hu-1 reference strain and an IC_50_ of 2.116 μM for the SARS-CoV-2 (D614G) variant, which indicated that teicoplanin inhibited authentic viruses at a relatively low and safe dose. Moreover, our data showed that the pre-treatment of teicoplanin was able to prevent authentic SARS-CoV-2 infection in hACE2 mice. Given that the principles of antiviral therapy are to prevent virus infection and use extensively as early as possible, it is reasonable to recommend the use of teicoplanin for SARS-CoV-2 in the early infection stage. Therefore, teicoplanin could potentially function as a dual inhibitor for both SARS-CoV-2 and co-infected Gram-positive bacteria.

## Materials and Methods

### Cell lines and viruses

HEK293T and Vero E6 cells were maintained in DMEM (ThermoFisher) supplemented with 10% FBS (ThermoFisher), 100 units/ml penicillin, and 100 μg/ml streptomycin (ThermoFisher) at 37 °C and 5% CO_2_. The HEK293T-hACE2 cell line was generated by infecting HEK293T cells with lentiviruses which expressed human angiotensin-converting enzyme 2 (hACE2). The hACE2-positive cells were sorted by fluorescence activated cell sorting (FACS) and confirmed by western blot with antibodies against hACE2. The HEK293T-hACE2 cells were maintained as wildtype HEK293T cells. All cells have been tested for mycoplasma by PCR-based assay and confirmed to be mycoplasma-free (Mycoplasma-F: 5’-GGGAGCAAACAGGATTAGTATCCCT-3’; Mycoplasma-R:5’-TGCACCATCTGTCACTCTGTTACCCTC-3’).

The plasmid expressing the Spike (S) of SARS-CoV-2 (D614; Wuhan-Hu-1, GISAID: EPI_ISL_402125) was purchased from Generay Biotech company (Shanghai, China) and inserted into the pcDNA3.1 vector. SARS-CoV-2 S / HIV-1 pseudotyped viruses were packaged by co-transfecting a lentiviral construct pHIV-Luciferase (Addgene plasmid # 21375), a packaging construct psPAX2 (Addgene plasmid # 12260) and a plasmid expressing S proteins into HEK293T cells. The culture medium was replaced with fresh DMEM 6 hours post transfection. The pseudotyped viruses-containing supernatant was collected 48 hours post transfection and filtered through 0.45 μm filters. The amounts of pseudotyped viruses were quantified by RT-qPCR assay with primers against the long term repeat (LTR) (HIVTotal-F: 5’-CTGGCTAACTAGGGAACCCACTGCT-3’; HIVTotal-R: 5’-GCTTCAGCAAGCCGAGTCCTGCGTC-3’). Pseudotyped viruses including SARS-CoV S / HIV-1 and VSV-G / HIV-1 were packaged and quantified as pseudotyped SARS-CoV-2 S / HIV-1 viruses. Different pseudotyped SARS-CoV-2 S / HIV-1 mutants were also packaged and quantified as above, the S proteins of which included those of G614 virus (SYSU-IHV, EPI_ISL_444969), B.1.1.7 (Alpha, GISAID: EPI_ISL_581117), B.1.351 (Beta, EPI_ISL_678597), P.1 (Gamma, EPI_ISL_792683), B.1.429 (Epsilon, EPI_ISL_1675148), B.1.525 (Eta, EPI_ISL_1093465), B.1.526 (Iota, EPI_ISL_1080752), B.1.617.1 (Kappa, EPI_ISL_1372093), B.1.617.2 (Delta, EPI_ISL_1337507), B.1.621 (Mu, EPI_ISL_1220045) and C.37 (Lambda, EPI_ISL_1534645).

Patient-derived authentic SARS-CoV-2 D614 virus (Wuhan-Hu-1) was obtained from Guangdong Provincial Center for Disease Control and Prevention. Authentic SARS-CoV-2 G614 virus (SYSU-IHV) was isolated from the sputum sample of a female admitted in Guangzhou Eighth People’s Hospital who was infected at Guangzhou by an African traveler in April 2020. Vero E6 cells were utilized to propagate these viruses.

### Pseudotyped virus infection assay

Pseudotyped viruses including SARS-CoV S / HIV-1, SARS-CoV-2 S / HIV-1, VSV-G / HIV-1 and 10 different SARS-CoV-2 mutants were packaged as above. For viral infection in siRNA-mediated gene knock-down experiment, targeted genes in HEK293T cells were firstly knocked down by a mixture of three different siRNAs, followed by the infection of pseudotyped SARS-CoV-2 S / HIV-1 virus 24 hours post transfection. Another 24 hours later, siRNA- and virus-treated cells were lysed and measured for the amounts of luciferase which could represented the percentages of virus-infected cells.

For virus infection in protein overexpression experiment, HEK293T cells were firstly transfected with CTSL-expressing or hACE2-expressing plasmids, followed by infecting with pseudotyped SARS-CoV-2 S / HIV-1 virus. The amounts of luciferase within each group were measured utilizing luminometer (Promega) 48 hours post infection, and represented as relative luminescence units.

For virus infection in drug treatment experiment, HEK293T-hACE2 cells were incubated with serially diluted drugs including teicoplanin (Selleck, S1399), dalbavancin (Selleck, S4848) and vancomycin (Selleck, S2575), and different pseudotyped viruses. The amounts of luciferase within each group were measured 48 hours post infection.

### Authentic virus infection assay

HEK293T-hACE2 cells were seeded in 12-well plates. 24 hours post seeding, cells were co-incubated with authentic SARS-CoV-2 D614 (Wuhan-Hu-1) virus and two-fold serially diluted teicoplanin. Another 48 hours post incubation, the supernatant in each group was collected and proceeded to RNA extraction with RNeasy Mini Kit (QIAGEN, 74104) according to the manufacturer’s instruction. SARS-CoV-2 viral RNA copies were determined by one-step SARS-CoV-2 RNA detection kit (PCR-Fluorescence Probing) (Da An Gene Co., DA0931) with the following primers and probes: N-F (5’-CAGTAGGGGAACTTCTCCTGCT-3’), N-R (5’-CTTTGCTGCTGCTTGACAGA-3’) and N-P (5’-FAM-CTGGCAATGGCGGTGATGCTGC-BHQ1-3’). The half maximal inhibitory concentration (IC50) of teicoplanin against SARS-CoV-2 D614 virus was calculated by GraphPad software (San Diego, USA) according to these viral RNA copies within each group. The IC50 of teicoplanin against authentic SARS-CoV-2 G614 virus (SYSU-IHV) was determined similarly. Authentic SARS-CoV-2 virus infection assays were conducted in the BSL-3 facility of Sun Yat-sen University.

### Drug or virus pre-treatment assay

To determine whether teicoplanin targeted virus directly or targeted host cell indirectly, HEK293T-hACE2 cells were treated with drug or virus in three different ways. In the first group, HEK293T-hACE2 cells were pre-treated with 0 μM, 12.5 μM, 25 μM and 50 μM teicoplanin respectively. Four hours post drug treatment, cells were infected with pseudotyped SARS-CoV-2 S / HIV-1 virus. Another 48 hours later, the amounts of luciferase within each group were monitored and represented as relative luminescence units. In the second group, HEK293T-hACE2 cells were co-treated with different concentrations of teicoplanin and pseudotyped SARS-CoV-2 S / HIV-1 virus simultaneously. The amounts of luciferase were measured 48 hours post co-treatment. In the third group, cells were pre-infected with pseudotyped SARS-CoV-2 S / HIV-1 virus. Four hours post infection, cells were treated with different concentrations of teicoplanin. Another 48 hours later, cells were lysed and measured for the amounts of luciferase.

### *In vitro* CTSL enzymatic inhibition assay

To evaluate CTSL enzymatic activity upon teicoplanin treatment *in vitro*, the purified 250 ng CTSL proteins (Sino Biological Inc., 10486-H08H) were added into CTSL assay buffer (400 mM NaAc, 4 mM EDTA, 8 mM DTT and pH 5.5) and incubated in ice for 15 min for activation. To evaluate the inhibition of teicoplanin on CTSL activity, the purified 250 ng CTSL proteins were also co-incubated with 50 μM teicoplanin (Selleck, S1399). After activation, 2 μg *in vitro* purified SARS-CoV S (40634-V08B) or SARS-CoV-2 S (40589-V08B1) were added into each group. S protein only group (without CTSL and teicoplanin) was set as control group. The S-CTSL-teicoplanin mixtures were incubated at 37 °C for 1.5 hours. The enzymatic reaction was stopped by adding with SDS-PAGE loading buffer and followed by boiling at 100 °C for 10 min. Digested proteins were proceeded to SDS-PAGE and analyzed by silver staining (Sigma-Aldrich, PROTSIL2-1KT).

### Animal infection

Eight-week-old specific-pathogen-free (SPF) transgenic hACE2 mice (C57BL/6) were purchased from GemPharmatech Co., Ltd (Cat No.: T037657). All mice were housed in SPF facilities at Laboratory Animal Center of Sun Yat-sen University. Animal experiments were conducted in strict compliance with the guidelines and regulations of Laboratory Monitoring Committee of Guangdong Province of China. The Ethics Committees of Guangdong Provincial People’s Hospital and Sun Yat-sen University approved animal experiments.

Six hours before viral challenge, four hACE2 mice were intraperitoneally administrated with 100 mg/kg body weight teicoplanin (dissolved in saline). Four hACE2 mice in control group were intraperitoneally administrated with equal volume of saline. Six hours later, all the mice were intranasally challenged with 1×10^5^ focus-forming units (FFU) of authentic SARS-CoV-2 D614 virus. Another 5 days later, mice were euthanized to harvest lung tissues. Viral RNA copies in lung tissues were quantified by one-step SARS-CoV-2 RNA detection kit (PCR-Fluorescence Probing) (Da An Gene Co., DA0931). Authentic SARS-CoV-2 challenge studies were approved by the Ethics Committee of Zhongshan School of Medicine of Sun Yat-sen University on Laboratory Animal Care (Assurance Number: SYSU-IACUC-2021-B0020).

### Histopathology and immunohistochemistry

Lung tissues of authentic SARS-CoV-2 infected mice were fixed in 4% paraformaldehyde for at least two days. These lung tissues were embedded in paraffin and proceeded to histopathology and immunohistochemistry analysis (Nanjing FreeThinking Biotechnology Co., Ltd). For histopathology analysis, sections (3-4 μm) of lung tissues were stained with hematoxylin and eosin (H&E). For immunohistochemistry analysis, sections of lung tissues were deparaffinized and rehydrated with xylene and gradient alcohol. Antigens were retrieved in citric acid buffer (pH 6.0) and quenched with 3% H_2_O_2_. After blocking with BSA, sections were incubated with rabbit anti-SARS-CoV-2 Nucleoprotein (N) for 24 hours at 4 °C, followed by incubating with goat anti-rabbit IgG secondary antibody (HRP-conjugated) and staining with 3,3’-diaminobenzidine. Antibody-conjugated sections were stained with hematoxylin, followed by dehydrating with gradient ethanol. Samples were covered by neutral balsam and imaged with HS6 microscope (Sunny Optical Technology Co., Ltd).

### Sequence data collection and alignment

The genome sequences of SARS-CoV and SARS-CoV-2 were collected from the GenBank database (https://www.ncbi.nlm.nih.gov/labs/virus/vssi/#/) and the GISAID’s EpiCoV database (https://www.gisaid.org/). The sequences of SARS-CoV circulating in 2003 contain 6 strains (accession numbers: AY278488, AY545918, AY545917, AY394977, AY394978, and AY394979). The sequences of SARS-CoV-2 include 12 variants: D614 (Wuhan-Hu-1, GISAID: EPI_ISL_402125), G614 (SYSU-IHV, EPI_ISL_444969), B.1.1.7 (Alpha, GISAID: EPI_ISL_581117), B.1.351 (Beta, EPI_ISL_678597), P.1 (Gamma, EPI_ISL_792683), B.1.429 (Epsilon, EPI_ISL_1675148), B.1.525 (Eta, EPI_ISL_1093465), B.1.526 (Iota, EPI_ISL_1080752), B.1.617.1 (Kappa, EPI_ISL_1372093), B.1.617.2 (Delta, EPI_ISL_1337507), B.1.621 (Mu, EPI_ISL_1220045) and C.37 (Lambda, EPI_ISL_1534645). The S gene sequences were obtained from the genome of SARS-CoV and SARS-CoV-2 according to the annotation in the GenBank database. The sequence datasets were aligned using the ClustalW program implemented in MEGA X software. Consensus sequences were created using BioEdit software (http://www.mbio.ncsu.edu/bioedit/bioedit.html) based on the multiple alignment of SARS-CoV and SARS-CoV-2. The amino acid sequence logos were generated by WebLogo.

### Statistical analysis

All the measurements in this study have been performed for at least three times by at least two lab technicians or students. Detailed statistical information including statistical tests, sample numbers, mean values, standard errors of the mean (SEM) and p-values have been shown in the main text and figure legends. Statistical analysis was conducted with Graphpad Prism 8.0 or Microsoft Excel. Triplicate and quadruplicate data were presented as mean ± SEM. A value of p ≥ 0.05 was considered to be not statistically significant and represented as “ns”. A value of p < 0.05 was considered to be statistically significant and represented as asterisk (*). Value of p < 0.01 was considered to be more statistically significant and represented as double asterisks (**). Value of p < 0.001 was considered to be the most statistically significant and represented as triple asterisks (***). When comparing mean differences between groups which were split by one independent variable, one-way ANOVA with Tukey’s multiple comparison test or Dunnett’s multiple comparison test was conducted. When comparing mean differences between groups which were split by two independent variables, two-way ANOVA with Tukey’s multiple comparisons test or Dunnett’s multiple comparisons test was conducted. For data with a normal distribution, we used Student’s *t* test.

## Acknowledgements

This work was supported by National Natural Science Foundation of China (82102385) and National Postdoctoral Program for Innovative Talents of China Postdoctoral Science Foundation (BX20190398) to X.M. This work was also supported by the National Special Research Program of China for Important Infectious Diseases (2017ZX10202102 and 2018ZX10302103), the Special 2019-nCoV Project of the National Key Research and Development Program of China (2020YFC0841400), the Emergency Key Program of Guangzhou Laboratory (EKPG21-24), the Special 2019-nCoV Program of the Natural Science Foundation of China (NSFC) (82041002), the Special Research and Development Program of Guangzhou (202008070010), and the Important Key Program of NSFC (81730060) to H.Z. This work was also supported by National Natural Science Foundation of China (82102367) to F.Y. This work was also supported by National Natural Science Foundation of China (81971918), Shenzhen Science and Technology Program (Grant No. JSGG20200225150431472 and JCYJ20200109142601702), the Pearl River S&T Nova Program of Guangzhou (201806010118) and the Fundamental Research Funds for the Central Universities, Sun Yat-sen University (2021qntd43) to T.P. No potential conflict of interest was reported by the authors.

